# Definitely saw it coming? An ERP study on the role of article gender and definiteness in predictive processing

**DOI:** 10.1101/563783

**Authors:** Damien S. Fleur, Monique Flecken, Joost Rommers, Mante S. Nieuwland

## Abstract

People sometimes anticipate specific words during language comprehension. Consistent with word anticipation, pre-nominal articles elicit differential neural activity when they mismatch the gender of a predictable noun compared with when they match. However, the functional significance of this pre-nominal effect is unclear: Do people only predict the noun or do they predict the entire article-noun combination? We addressed this question in an event-related potential study (N=48) with pre-registered data acquisition and analyses, capitalizing on gender-marking on Dutch definite articles and the lack thereof on indefinite articles. Participants read mini-story contexts that strongly suggested either a definite or indefinite noun phrase (e.g., ‘het/een boek’, the/a book) as its best continuation, followed by a definite noun phrase with the expected noun or an unexpected, different gender noun (‘het boek/de roman’, the book/the novel). We observed an enhanced negativity (N400) for articles that were unexpectedly definite or mismatched the expected gender, with the former effect being strongest. Pre-registered analyses and exploratory Bayesian analyses did not yield convincing evidence that the effect of gender-mismatch depended on expected definiteness. While prediction of article form cannot be excluded, it may not be required to elicit pre-nominal effects.

## INTRODUCTION

Language comprehenders sometimes anticipate upcoming words based on the meaning of a story or conversation. Particularly informative in tracking the relevant anticipatory processes are event-related brain potentials (ERPs) recorded from the scalp. The ERP signal in response to words consists of various components including the N400 reflecting semantic processing (Kutas & Hillyard, 1980, 1984; for review, see Kutas & Federmeier, 2011) and post-N400 positivities in response to unexpected words that disconfirm likely expectations (for review, see Van Petten & Luka, 2012). Arguably the strongest evidence for word anticipation comes from pre-nominal manipulations, which measure behavioral or neural responses to an article or adjective appearing before a noun (for review, see Kutas, DeLong & Smith, 2011; Van Berkum, 2009). Most studies of this type use gender-marking of pre-nominal articles, such as in Spanish and Dutch, and report differential event-related potential (ERP) responses to articles that mismatch the gender of a highly predictable noun, compared to gender-matching articles (e.g., for Dutch, Otten and Van Berkum, 2009; Van Berkum et al., 2005; for Spanish, Foucart, Martin, Moreno & Costa, 2014; Gianelli & Molinaro, 2018; Martin, Branzi & Bar, 2018; Molinaro, Gianelle, Caffarra & Martin, 2017; Wicha, Bates, Moreno & Kutas, 2003; Wicha, Moreno & Kutas, 2003, 2004). Of particular relevance is a study by Otten and Van Berkum (2009), wherein participants read two-sentence mini-stories that contained an article-adjectives-noun combination of which the noun was either predictable (e.g., “de verfijnde maar toch opvallende <u>ketting</u>”, the_com_ sophisticated yet striking <u>necklace</u>_com_) or not predictable and of a different gender than the predictable noun (e.g., “het verfijnde maar toch opvallende <u>collier</u>”, the_neu_ sophisticated yet striking <u>collar</u>_neu_). The gender-mismatching articles elicited an N400-like differential ERP effect, which was not observed for the same article-adjective-noun combinations in non-constraining contexts featuring the same content words. Given that this was a comparison between words that were grammatical and did not differ in meaning, the observed effect must be ascribed to the grammatical relation between the pre-nominal word and the predicted - but not yet presented - noun.

Although the available literature supports noun prediction, the precise functional significance of pre-nominal effects remains unclear. A minimal interpretation is that people predict the noun (with or without its gender) and then use article gender, once available, to confirm or change the noun prediction (e.g., Otten & Van Berkum, 2009; see also Otten, Nieuwland & Van Berkum, 2007; Otten & Van Berkum, 2008; Van Berkum et al., 2005). However, a stronger claim has been made, namely that people predict a specific article-noun combination including the gender-marked form of the article (Kutas, DeLong & Smith, 2011; Wicha et al., 2003, 2004; DeLong et al., 2005; see also Dell & Chang, 2014; Van Petten & Luka, 2012). We contrasted these accounts in an ERP study on Dutch mini-story comprehension, with pre-registered data collection and analyses. The rationale of the experiment was that if the articles themselves are predicted, one would expect little effect of gender-match on articles in a situation where gender-marking was not expected in the first place. As will be explained later, Dutch provides such a situation.

In addition to these theoretical considerations, there is inconsistency in the type of effect that has been observed empirically. In the first published study with a pre-nominal gender manipulation (Wicha et al., 2003), Spanish speakers listened to sentence pairs in which a predictable noun or an incongruent noun was replaced with a drawing. The authors observed greater N400 amplitude for gender-marked pre-nominal articles that mismatched the gender of the predictable nouns, compared to articles with matching gender. A follow-up study with written materials and line drawings (Wicha et al., 2003) also obtained an N400 effect of gender-mismatch. In a follow-up with fully spoken sentences and no line drawings (Wicha et al., 2004), gender-mismatching articles now elicited a P600 effect, which was interpreted as indicating an article-noun agreement violation. Recent studies with written Spanish sentences show predominantly N400-like effects in relation to gender-mismatching articles, i.e. enhanced negativities in the typical N400 time window^1^ (Foucart et al., 2014; Martin et al., 2018; Molinaro et al., 2014).

The first Dutch study on prediction did not use pre-nominal articles but adjectives (Van Berkum et al., 2005). The participants listened to mini-stories that contained either a highly predictable noun, or a different-gender, non-predictable noun. The nouns were preceded by adjectives that were gender-marked (using the adjectival suffix rule that adds ‘-e’ to neuter nouns) in agreement with the upcoming noun. Time-locked to inflection-onset, gender-mismatches elicited an early positivity between 50 and 250 ms compared to gender-matches. Two follow-up studies with the same manipulation (Otten et al., 2007; Otten & Van Berkum, 2008) reported different ERP effects. In a study with spoken stories (Otten et al., 2007), gender-mismatching adjectives elicited a negative, right-frontal ERP effect between 300 and 600 ms after adjective-onset. In a study with written materials (Otten & Van Berkum, 2008), gender-mismatches elicited a late negativity ERP effect at 900-1200 ms after adjective-onset. In the Otten and Van Berkum (2009) study discussed previously, a negativity was observed in the 200-600 ms time window at right-frontal electrodes, which grew in size over time. Finally, Kochari & Flecken (2018), using the Otten and Van Berkum materials but omitting the non-constraining contexts, did not obtain a statistically significant effect of gender-mismatch. Mismatching articles did elicit a slowly developing negative shift compared to matching articles, over posterior electrodes instead of frontal electrodes. The observed pattern was consistent with that in the original data in terms of effect size (leaving aside differences in scalp-distribution), but a Bayesian analysis supported neither the null-hypothesis (no prediction effect) nor the alternative hypothesis (the effect size reported by Otten and Van Berkum).

### Three open questions

The current study aimed to shed light on the following three questions. First, do people only predict a noun, and then use the information that the article provides to update their prediction, or do they predict the specific gender-marked article *itself*, along with the meaning and form of the noun? On the first account, the initial prediction could be limited to a specific noun meaning (either without activation of gender information or with activation of gender^2^ but not of an article form, which depends on gender and definiteness). Once the article is presented, the available gender information can be used to update the prediction. Whether or not noun gender was activated before the article appeared, the processes elicited by the article are also predictive because they involve access to the gender of the yet unseen noun. On the second account (DeLong et al., 2005; Kutas et al., 2011; Wicha et al. 2003, 2004; for discussion, see Ito et al., 2017c), gender and definiteness information is activated before the article appears, for example as part of a lexical pre-activation process where people access syntactic and semantic information associated with a specific word form. The main difference between these accounts, therefore, is whether or not people predict a specific article word form (i.e., a lexical prediction).

Observing gender-mismatch effects on words that are not articles, such as adjectives (Van Berkum et al., 2005) suggests that prediction of a pre-nominal word-form is not required to elicit an effect. However, those effects are not consistent across studies and do not seem to involve modulation of the N400 (for discussion, Ito et al., 2017a,c), which leaves open the possibility that pre-nominal N400 effects do reflect lexical form prediction. Previously reported effects of article gender-mismatch are in principle consistent with both these accounts. Some authors explicitly advocate article form prediction (Wicha et al., 2004, 2004; see also DeLong et al., 2005), whereas others have only advocated noun prediction (e.g., Otten et al., 2007; Otten & Van Berkum, 2008, 2009; Van Berkum et al., 2005). In the current study, we therefore tried to tease apart these accounts by testing for gender-mismatch effects on articles that *themselves* are either expected or unexpected in terms of another feature, i.e., definiteness. We capitalized on the fact that in Dutch, like in other gender-marking languages such as Spanish and German, the specific article form depends not only on gender but also on definiteness. Dutch definite articles are gender-marked (‘de’ for common-gender nouns, ‘het’ for neuter-gender nouns), whereas indefinite articles are not (‘een’ for nouns of either gender).

Our second question is: what is the role of article definiteness during predictive processing? In languages that mark both gender and definiteness on the article (e.g., Dutch and Spanish), the article contains lexical information and semantic/referential information that is relevant to interpretation (e.g., Abbott, 2004, 2006; Frazier, 2006; Heim, 1982). Previous experiments on Spanish have compared gender-matching and -mismatching articles that are both either definite or indefinite. In Dutch, however, definite articles are gender-marked while indefinite articles are not, which is why Otten and Van Berkum (2009) and Kochari and Flecken (2018) only used definite articles. Both the Spanish and the Dutch studies used the same sentence completion procedure (‘cloze test’) to establish predictability, in which participants completed sentences truncated before the article. But while the Spanish cloze values directly reflected the obtained article-noun responses, the Dutch cloze values only reflected the noun responses, and the gender-manipulation with definite articles was implemented for sentence contexts where most completions involve an indefinite article, at least for some items^3^. Therefore, some of the definite articles in Otten & Van Berkum (2009) and Kochari & Flecken (2018) were probably unexpected or infelicitous regardless of their gender. The contexts that license the introduction of a novel definite referent are more limited or restricted than those that license the introduction of novel indefinite reference (for discussion, see Abbott, 2004, 2006; Clifton, 2013; Fraurud, 1990, Frazier, 2006; Heim, 1982; Singh, Fedorenko, Mahowald, & Gibson, 2016), and definite reference is more commonly used for previously mentioned referents than for new referents. Unexpected or infelicitous definiteness of the article may itself incur a processing cost reflected in N400 amplitude (Kirsten, Tiemann, Seibold, Hertrich, Beck & Rolke, 2014; Schlueter, Namyst & Lau, 2018; see also Anderson & Holcomb, 2005; Schumacher, 2009). This could indicate that (in)definiteness conveys meaning and therefore results in additional semantic processing, or even that people predict the definiteness of upcoming referents or have difficulty integrating the article into an event-based representation of the discourse context (a ‘situation model’; Zwaan et al., 1995). The results of Otten and Van Berkum (2009) and Kochari and Flecken (2018) thus reflect an unknown mix of effects associated with gender and definiteness, meaning that it is unclear whether people in fact predict pre-nominal lexical material. The current study, therefore, tested for gender-mismatch effects on articles that are either expectedly or unexpectedly definite.

Finally, examining these issues in Dutch could provide insights into the consistency of article gender-mismatch effects across languages. There is qualitative variability in the observed ERP effects of gender mismatch. This variability may signal something meaningful like cross-linguistic differences or differences associated with specific methodological choices, it may signal random fluctuations (noise), or an unknown mix of the above (for discussion, see Ito et al., 2017c). Almost all the studies with Romance languages such as Spanish, Catalan or Italian report N400 effects (e.g., Wicha et al., 2003; Foucart et al., 2014; Martin et al., 2018; Molinaro et al., 2014). There is only one Spanish study reporting a P600 effect (Wicha et al., 2004), which, to our knowledge, has not yet been replicated. In addition, there are two Dutch studies reporting ‘N400-like’ effects with different scalp distributions (Kochari & Flecken, 2018; Otten & Van Berkum, 2009), which are most relevant to the current study. We believe there is reason to doubt that the patterns observed in these two Dutch studies are truly generated by the article^4^. It is currently unclear why Kochari and Flecken (2018) and Otten and Van Berkum (2009) report different ERP results, and why these effects differed from effects reported for other languages, mainly Romance languages in which article gender is arguably less ambiguous: the Dutch definite articles ‘de’ and ‘het’, besides signalling a singular noun of common or neuter gender, can also signal plurals and diminutives, irrespective of gender. The effect of gender-mismatch in Dutch itself may have been diluted by items where the article itself was unexpectedly definite. This would not have occurred in the studies with languages like Spanish or Italian, which have gender-marking on definite and indefinite articles and separate marking for plurality. By controlling the expectedness of the articles in the current study, we tried to get a better view on the type of effect elicited by gender-mismatch, in Dutch specifically.

### The current study

In the current ERP study, we investigated lexical prediction during Dutch mini-story comprehension, and we addressed three outstanding questions on ERP effects associated with a gender-mismatch between a pre-nominal article and a predictable noun. We asked (1) whether such effects occur for predictable and unpredictable articles alike, (2) whether article definiteness mismatch has an effect alongside gender mismatch, and (3) whether a gender-mismatch effect in Dutch elicits enhanced N400 amplitude like in other languages if definiteness match is controlled for.

Our participants read two-sentence mini-stories in four different conditions (see Table 1 for an example item), with the critical noun phrase embedded in the second sentence. Each participant read one of two contexts that suggested a specific noun as its best continuation as part of either a definite noun phrase or an indefinite noun phrase (as established in a cloze task, see Methods). Each context was followed by a definite noun phrase, which either contained the predictable noun or an unpredictable, different-gender noun. Because half of the stories contained unexpectedly definite articles, we included filler stories with predictable indefinite noun phrases, such that a mismatch of the expected definiteness was as likely as a mismatch of the expected gender.

**Table 1.** Dutch example mini-story in each of the four conditions, plus approximate translation. The entire set of materials is available on osf.io/6drcy

|  |  |  |  |
| --- | --- | --- | --- |
|  | Context | Critical noun phrase | Sentence ending |
| Expectedly definite | *Het is zondagochtend. De gehele gelovige familie gaat zoals altijd naar* | **Gender-match:**<br>*de<sub>com</sub> kerk<sub>com</sub>*<br>**Gender-mismatch:**<br>*het<sub>neu</sub> gebedshuis<sub>neu</sub>* | *in het dorp.* |
| Unexpectedly definite | *Mijn moeder is erg gelovig. Op vakantie gaan we altijd direct op zoek naar* | **Gender-match:**<br>*de<sub>com</sub> kerk<sub>com</sub>*<br>**Gender-mismatch:**<br>*het<sub>neu</sub> gebedshuis<sub>neu</sub>* | *in de stad.* |
| Approximate translation |  |  |  |
| Expectedly definite | *It is sunday morning. The whole religious family goes, as always, to* | **Gender-match:**<br>*the<sub>com</sub> church<sub>com</sub>*<br>**Gender-mismatch:**<br>*the<sub>neu</sub> worship place<sub>neu</sub>* | *in the village.* |
| Unexpectedly definite | *My mother is really religious. When on vacation, we always look directly for* | **Gender-match:**<br>*the<sub>com</sub> church<sub>com</sub>*<br>**Gender-mismatch:**<br>*the<sub>neu</sub> worship place<sub>neu</sub>* | *in the city.* |

Our study was not a direct replication attempt of Otten and Van Berkum (2009), nor of Kochari and Flecken (2018). Our experimental design was different, and we pre-registered different analyses. Our primary dependent variable was N400 amplitude, defined as the average voltage value in the 300-500 ms time window after word onset at a centroparietal electrode selection. We defined additional dependent variables for anterior electrodes and for the subsequent time-window (500-700 ms) to capture later activity like extended N400 effects, the PNP or P600 (DeLong, Quante, & Kutas, 2014; Nieuwland et al., 2019; Van Petten & Luka, 2012). We predicted that article gender-mismatch would elicit enhanced N400 amplitude compared to gender-match, like the patterns observed in Spanish and Italian (Wicha et al., 2003; Foucart et al., 2014; Martin et al., 2018; Molinaro et al). This would be consistent with pre-activation of the noun, but would not suffice to conclude participants predicted article form. In addition, we predicted that unexpectedly definite articles would elicit enhanced N400 amplitude compared to expectedly definite articles (e.g., Schlueter et al., 2018).

Our central question was whether or not we would observe an interaction pattern. If gender-mismatch effects reflect the prediction of specific article-noun combinations, including article form (DeLong et al., 2005; Wicha et al., 2004), then we expect to observe an interaction effect: a gender-mismatch effect for expectedly definite articles but not for unexpectedly definite articles. Alternatively, if gender-mismatch effects only reflect the incremental use of article gender-information to update a prediction, rather than the consequences of an article form misprediction, we expect to observe no interaction. We considered a third, hybrid option wherein people predict specific articles but also use gender information on unpredicted articles to update their prediction, which would be supported by a gender-mismatch effect that was obtained for both expectedly and unexpectedly definite articles but that was larger for expectedly definite articles. Finally, we considered a fourth option, wherein the effect of gender-mismatch is qualitatively different for expectedly and unexpectedly definite articles (e.g., a P600 effect for expectedly definite articles and a N400 effect for unexpectedly definite articles, or vice versa), which would support a processing distinction between the detection of an article prediction-mismatch and the incremental use of gender-information during predictive processing.

## METHODS

### Participants

We recruited 48 participants^5^ (17 males; participant gender was not pre-registered; mean age of 24 years, range 19-33) from the participant pool of the Max Planck Institute for Psycholinguistics in Nijmegen, the Netherlands. We did not perform an a priori power analysis to determine the required sample size, but we decided on a sample size that was a multiple of 4 and larger than previous studies at the time. All participants were native Dutch speakers, right-handed, with normal or corrected-to-normal vision and without history of language impairment. After receiving information about the experimental procedures, participants gave informed written consent to take part in the experiment, which was approved by the Ethics Committee for Behavioural Research of the Social Sciences Faculty at Radboud University Nijmegen in compliance with the Declaration of Helsinki. Participants were paid for their participation (18 €). Participant data were excluded from further analysis based on pre-registered criteria about the number of artefact-free trials (fewer than 25 trials in any of the conditions, or fewer than 30 trials on average across conditions) and the accuracy with which they answered the comprehension questions (<80% correct). We excluded and replaced 3 additional participants to achieve our pre-registered sample size.

### Materials

The final set of materials for this study was selected from a larger set based on specific constraints. We initially created a set of 280 items, of which each item contained two different versions of a Dutch mini-story. The two-sentence stories were written such that one version presumably led people to expect a specific definite noun phrase (the ‘definite context’), and the other version presumably led people to expect that same noun as part of an indefinite noun phrase (the ‘indefinite context’). The definite and indefinite contexts sometimes differed in the number of words in the first sentence, but always contained the same number of words in the second sentence (i.e., the sentence position of the target words was matched between versions, but not between items). To establish whether the stories indeed were sufficiently constraining towards these noun phrases, we performed a cloze probability test in the form of an online questionnaire. All mini-stories were truncated before the target article. We created two lists of 280 stories such that each participant saw only one version of each item. Within each list, we randomized definite and indefinite contexts. We recruited 40 participants (20 per list) from the pool of participants of the Max Planck Institute of Psycholinguistics in Nijmegen who received a financial compensation (10€). They were instructed to read each mini-story in the order they were given, and complete each item with the continuation they would have expected. Participants were given an example of a mini-story with a possible ending that matched the structure of the test items. They received no specific instruction regarding the number of words to use but were asked to avoid repeating words over multiple stories, to not think too long about a specific story and to use whatever completion came to mind first.

From the obtained responses, we counted how often the expected article and/or noun was used. We also counted certain answers towards the target noun when the response had lexical overlap with and the same gender as the target noun (e.g. ‘de pc’ for ‘de computer’), when the response was a misspelling or differently-spelled version of the target noun, when the response was the diminutive version of a neuter-gender target noun (e.g. ‘het spelletje’ for ‘het spel’). Cloze probability was calculated as the percentage of responses containing the target article or target noun. We selected the items in which each version had a cloze probability of at least 75% for the definite and indefinite target article and the noun, and where an unexpected article of the wrong gender or definiteness was never higher than 15%. For 89 items that did not make this selection, we rewrote one or both versions and performed a second cloze test with 20 participants who had not participated in the first cloze test, and we computed new cloze probability scores and again selected items that made the 75% cloze probability cut-off for both articles and the noun.

The final selection contained 160 items, with an average cloze value for the expected target article of 94% (SD = 7, range = 75-100) and 92% for the expected target noun (SD = 8, range = 75-100). Of note, gender was not fully balanced across items, because we had 99 target nouns of common gender (de-words) and only 61 of neuter gender (het-words). This disparity matches the relatively high frequency of de-words compared to het-words (Deutsch & Wijnen, 1985; Tuinman, 1996, quoted in Geerts, 1975; Van Berkum, 1997). We controlled for a potential effect of the article form in our statistical analyses described below. On average, the target article was the 8th word in the second sentence (SD = 1.9, range = 3-13) and the target noun followed right after. Sentence position of the target article and noun was matched for the definite and indefinite context of each item.

For the ERP experiment, we created the gender-mismatch condition by replacing the target article-noun combination with an unexpected, different-gender article-noun combination (Article: Mean = 1%, SD = 3, range = 0-15; Noun: Mean = 0%, SD = 2, range = 0-20). We selected mismatching nouns that we considered relevant and at least somewhat plausible or non-anomalous given the story context. To create the unexpectedly definite condition, we then replaced the expected indefinite article (‘een’) of each indefinite-context with a definite article of the correct gender. In addition, we added at least one and at most three words after the target noun, and this sentence-ending was identical for the definite and indefinite context.

In the experimental stories, an expectation of an indefinite noun phrase was never met. To avoid that participants would pick up on this regularity (and therefore, possibly, would stop predicting indefinite noun phrases), we included 80 filler stories with a high-cloze indefinite noun phrase (Article: Mean = 94%, SD = 3, range = 75-100%; Noun: Mean = 91%, SD = 8, range = 75-100). The fillers were generated from the set of materials that did not make it into the experimental materials, and were of the same two-sentence form as our experimental materials (e.g., “*Lisa’s dochter lijkt koorts te hebben. Om de temperatuur te meten leent ze bij de buurvrouw EEN thermometer voor kinderen”*, approximate translation: ‘Lisa’s daughter seems to have a fever. To measure the temperature, she borrows from the neighbour A thermometer for children’, critical article capitalized for demonstration purpose only). Due to the fillers, ERP participants saw the same ratio of unexpectedly definite articles and articles with an unexpected gender compared to expected articles, namely in ⅓ of all stories.

In our experimental materials, we manipulated the variables expected article definiteness and article gender-match in a 2 (Definiteness: expected, unexpected) by 2 (Gender: mach, mismatch) factorial design. We created 4 stimulus lists such that each participant saw 40 items from each of the 4 conditions, and each participant saw only one condition of an item, but across the lists each item was seen in each condition equally often. For each stimulus list, we generated two randomizations, to a total of 8 lists.

To encourage participants to pay attention to the meaning of the stories, they were asked to answer yes/no comprehension questions on 60 trials (i.e., 25% of all trials were followed by a question). These comprehension questions were roughly evenly spread across the experiment and separated from each other by at least two trials.

### Procedure

Participants were seated before a monitor in a soundproof, electrically shielded room. Using a button box, participants could start each trial, which started with a fixation cross displayed at the centre of the screen, followed by the first sentence of a story shown in its entirety. Participants could press a button to start the second sentence, which was presented one word at a time at the centre of the screen. Word duration was 300 ms and was followed by a blank screen for 300 ms until the next appeared. If the story was followed by a comprehension question, participants were required to respond yes or no with the button box before the next trial started.

A brief practice session with five trials preceded the actual experiment, so that participants could get used to the procedure. The experiment was divided in six blocks with brief breaks in between.

### EEG recording and data-processing

We recorded continuous EEG signal from 27 active scalp electrodes mounted in an elastic cap (ActiCap), placed according to the 10-20 convention and each referenced online to the left mastoid. An additional reference electrode was placed at the right mastoid. Furthermore, we recorded voltage at 4 EOG electrodes (above and under the left eye for the vertical dimension, next to the left and right eye for the horizontal dimension). The signal was amplified using BrainAmps amplifiers and recorded with Brain Vision Recorder (Brain Products, München) at 500 Hz, with a band-pass filter at 0.016-150 Hz (time constant 10s).

We used BrainVision Analyzer for offline data processing. Following the pre-registration, we visually screened the data for bad channels (due to drifting, spiking, excessive line noise) and interpolated bad channels through spline interpolation. We then filtered the continuous data with a 0.1-100 Hz (24 dB/octave roll-off) band-pass filter, and we re-referenced all channels to the average of the left and right mastoid. We then epoched the data into segments from −500 to 1000 ms relative to target article or noun onset. We subsequently removed artefact-containing segments (i.e., containing large movement-related artefacts, large bursts of muscle activity, or amplifier blocking) after visual inspection. We then performed an ICA-based correction for blinks, eye movements, and steady muscle activity. After this, we applied a 30 Hz low-pass filter (24 dB), followed by a baseline correction to 200 ms before each critical word. Finally, we automatically rejected segments with values that exceed ±75 µV at any channel. In total, 4.4% of the epoched data was removed.

### Statistical analyses

We performed linear mixed-effects analyses in R (version 3.3.3; R Core Team, 2018) with the R package ‘lme4’ (Bates, Maechler, Bolker & Walker, 2014), with the two-level factors ‘definiteness’ (expected/unexpected) and ‘gender’ (match/mismatch). Definiteness refers to the match between the definite article with the story context, as the articles were either expectedly definite or unexpectedly definite (high cloze values for definite or indefinite articles, respectively). Gender refers to whether the article matches the gender of the expected noun. We included an additional factor ‘article’ (de/het) to account for potential effects associated with the specific articles, which was important given the lexical differences between ‘de’ and ‘het’ (‘de’ is more frequent, and may elicit smaller N400s overall; Kutas & Federmeier, 2011), and given that most of our items had ‘de’ as the expectedly definite article.

Using a spatiotemporal region-of-interest (ROI) approach, our main dependent measure (N400 amplitude) was the average voltage across five centro-parietal channels (Cz, CP1, CP2, P3, Pz, P4) in the 300–500 ms window after word onset for each trial. To evaluate effects at anterior electrodes, we also computed average voltage across five anterior electrodes (F3, Fz, F4, FCz, FC1, FC2). For both ROIs, we also computed average voltage in the 500-700 ms (post-N400) time window. For the articles, we pre-registered analyses at both time windows in both ROIs, resulting in four analyses. For the nouns, we only pre-registered two analyses, namely on voltage in the 300-500 ms time window at the posterior ROI and the 500-700 ms time window at the anterior ROI. We deviation-coded the variables ‘definiteness’ and ‘gender’, and evaluated their effects by performing model-comparison using chi-square goodness-of-fit tests.

Following the recommendations of Barr, Levy, Scheepers, & Tily (2013), we first tried the maximal random effect structure as justified by the design but simplified the random effect structure to deal with non-convergence. For the article and noun analyses, we included random effects for subjects and items and by-subject and by-item random slopes for ‘gender’.

## RESULTS

### Pre-registered article-analyses

In accordance with our predictions, our experimental manipulations were associated with modulations of N400 activity, visible at posterior electrodes within the 300-500 ms time window after article onset (Figure 1; supplementary figures that show all channels can be found at https://osf.io/m9weg/). Our pre-registered analysis yielded the following patterns (see Table 2, for details): Gender-mismatching articles elicited reliably more negative voltage (enhanced N400 activity) compared to gender-matching articles at the posterior ROI. This effect extended into the 500-700 ms time window^6^, as also observed in previous studies with Spanish sentences (Martin et al., 2013, 2018; Foucart et al., 2014). Unexpectedly definite articles elicited more negative ERPs than expectedly definite articles at both ROIs and in both time windows, although this effect was strongest at the posterior ROI in the N400 time window, thus consistent with an N400 effect. The gender-mismatch effect was numerically larger for expectedly definite articles (−0.65 µV, *SE =* 0.27) than for unexpectedly definite articles (0.38 µV, *SE* = 0.26), but the results did not allow us to reject the hypothesis that these effect are similar. In keeping with our pre-registration protocol, we did not follow-up on this non-significant interaction pattern. In the 500-700 ms time window, where we also obtained effects of gender and definiteness, there was no hint of an interaction pattern because the estimate for the interaction term was close to zero. Figure 2 shows the scalp distribution of the article effects.

**Table 2.** Results from the pre-registered article-analyses. For each spatial and temporal region-of-interest, the tables shows the estimated difference between the expected and unexpected conditions (unexpected minus expected), the associated 95% confidence interval, the □^2^ test-result and associated p-value (for details, see analysis files on https://osf.io/6drcy).

|  |  | Time window |  |  |  |  |  |
| --- | --- | --- | --- | --- | --- | --- | --- |
| Factor | ROI | 300-500 ms<br>(N400) |  |  | 500-700 ms<br>(post-N400) |  |  |
| | | $\beta$, *CI* | $\chi^2$ | *p* | $\beta$, *CI* | $\chi^2$ | *p* |
| Gender | Anterior | -0.11, [-0.52, 0.30] | 0.29 | 0.59 | -0.18, [-0.62, 0.25] | 0.69 | 0.40 |
|  | Posterior | -0.52, [-0.94, -0.11] | 6.10 | 0.01 | -0.59, [-1.02, -0.17] | 7.20 | 0.01 |
| Definiteness | Anterior | -1.06, [-1.43, -0.69] | 31.07 | <0.001 | -0.42, [-0.83, -0.01] | 4.04 | 0.04 |
|  | Posterior | -1.19, [-1.55, -0.83] | 42.42 | <0.001 | -0.77, [-1.16, -0.39] | 15.43 | <0.001 |
| Gender :Definiteness | Anterior | 0.04, [-0.78, 0.71] | 0.01 | 0.92 | 0.02, [-0.80, 0.84] | 0.00 | 0.97 |
|  | Posterior | 0.40, [-0.31, 1.12] | 1.22 | 0.27 | 0.00, [-0.77, 0.77] | 0.00 | 1.00 |

**Figure 1.**
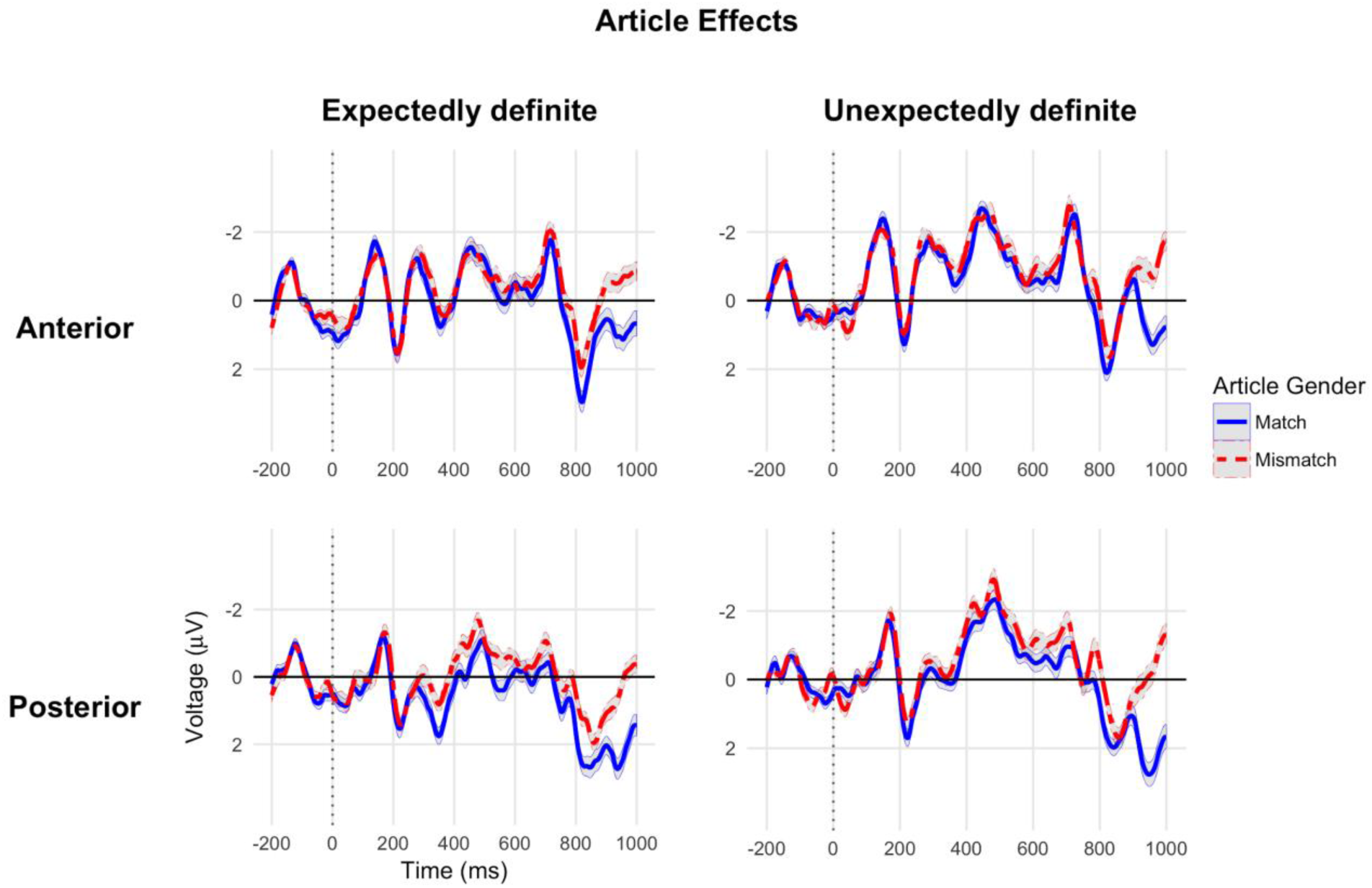
Article effects. The graphs show the grand-average ERPs elicited by gender-matching articles (solid blue lines) and gender-mismatching articles (dotted red lines) at the anterior and posterior ROIs (top and bottom graphs, respectively), when articles were expectedly and unexpectedly definite (left and right graphs, respectively). Grey-shaded areas show the within-subject standard error of the condition mean (Cousineau, 2005; Morey, 2008; calculated with the ‘Rmisc’ package in R). We emphasize that these ERP plots do not directly correspond to the results of our statistical analyses, which used linear mixed-effects to account for variance associated with different items and the two article forms (‘de/het’). Note that the large differences towards the end of the epoch reflect activity after noun onset.

**Figure 2.**
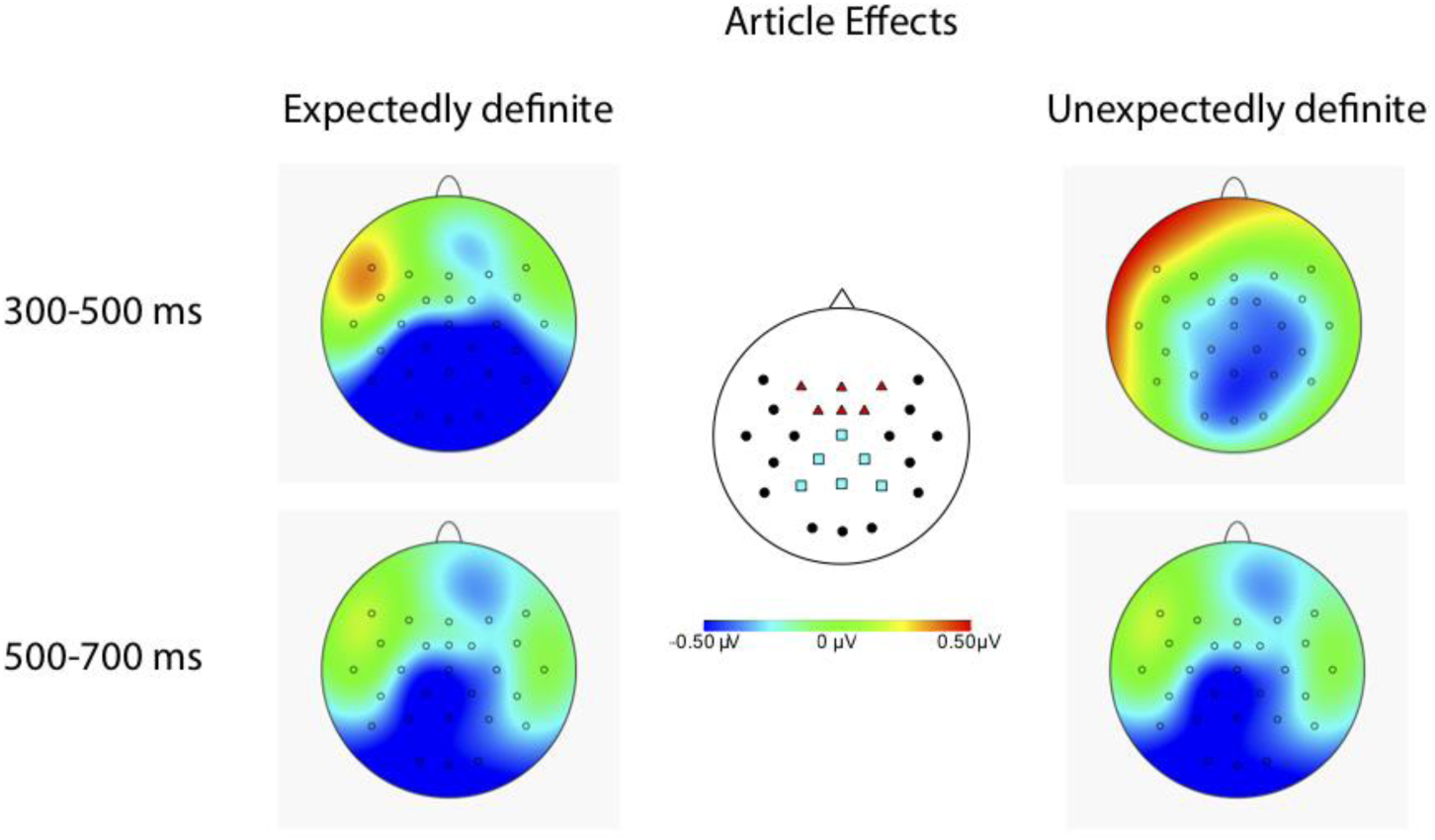
Scalp plots of the gender effects (mismatch minus match) for expectedly and unexpectedly definite articles in both time windows of analysis. Blue squares and red triangles in the center head plot show the positions of the electrodes contained in the pre-registered posterior and frontal ROI, respectively.

### Pre-registered noun-analyses

As expected, prediction-mismatching nouns elicited more negative ERPs in the posterior ROI at 300-500 ms after noun onset, i.e., an N400 effect, compared to matching nouns (Figure 3; Table 3), and more positive ERPs in the anterior ROI at 500-700 ms, although this later positive ERP effect appeared much weaker than the earlier N400 effect. In the posterior ROI at 300-500 ms, nouns following expectedly definite articles elicit more negative ERPs compared to nouns following unexpectedly definite articles. Finally, for the posterior ROI at 300-500 ms, but not for the anterior ROI at 500-700 ms, we found evidence against the same effect of mismatch regardless of definiteness; the N400 effect of prediction mismatch was more pronounced for unexpectedly definite nouns, ß = −1.45, CI = [−2.21, −0.68], □^2^(1) = 12.91, p < 0.001, than for expectedly definite nouns, ß = −0.82, CI = [−1.48, −0.16], □^2^(1) = 5.83, p = 0.01. The average voltages were less positive overall for nouns after expectedly definite articles (match, mean = 1.47 µV, *SE* = 0.41; mismatch, mean = 0.65 µV, *SE* = 0.33) than after unexpectedly definite articles (match, mean = 2.57 µV, *SE* = 0.40; mismatch, mean = 1.12 µV, *SE* = 0.38). The interaction pattern thus mostly resulted from the effect of definiteness on the matching nouns. Figure 4 shows that the N400 effect of prediction mismatch after expectedly definite articles had a slightly unusual frontal distribution. We will briefly return to the unanticipated patterns in the Discussion.

**Table 3.** Results from the pre-registered noun-analyses. For each spatial and temporal region-of-interest (ROI), the tables shows the estimated difference between the expected and unexpected conditions (unexpected minus expected), the associated 95% confidence interval, the □^2^ test-result and associated p-value.

|  | ROI |  |  |  |  |  |
| --- | --- | --- | --- | --- | --- | --- |
| Factor | Posterior, 300-500 ms |  |  | Anterior, 500-700 ms |  |  |
| | $\beta$, *CI* | $\chi^2$ | *p* | $\beta$, *CI* | $\chi^2$ | *p* |
| Mismatch | -1.14, [-1.64, -0.64] | 17.96 | <0.001 | 0.55, [0.01, 1.09] | 4.03 | 0.04 |
| Definiteness | 0.66, [0.27, 1.04] | 11.23 | <0.001 | 0.05, [-0.36, 0.45] | 0.06 | 0.81 |
| Mismatch:Definiteness | -0.87, [-1.64, -0.10] | 4.96 | 0.03 | 0.09, [-0.72, 0.90] | 0.05 | 0.83 |

**Figure 3.**
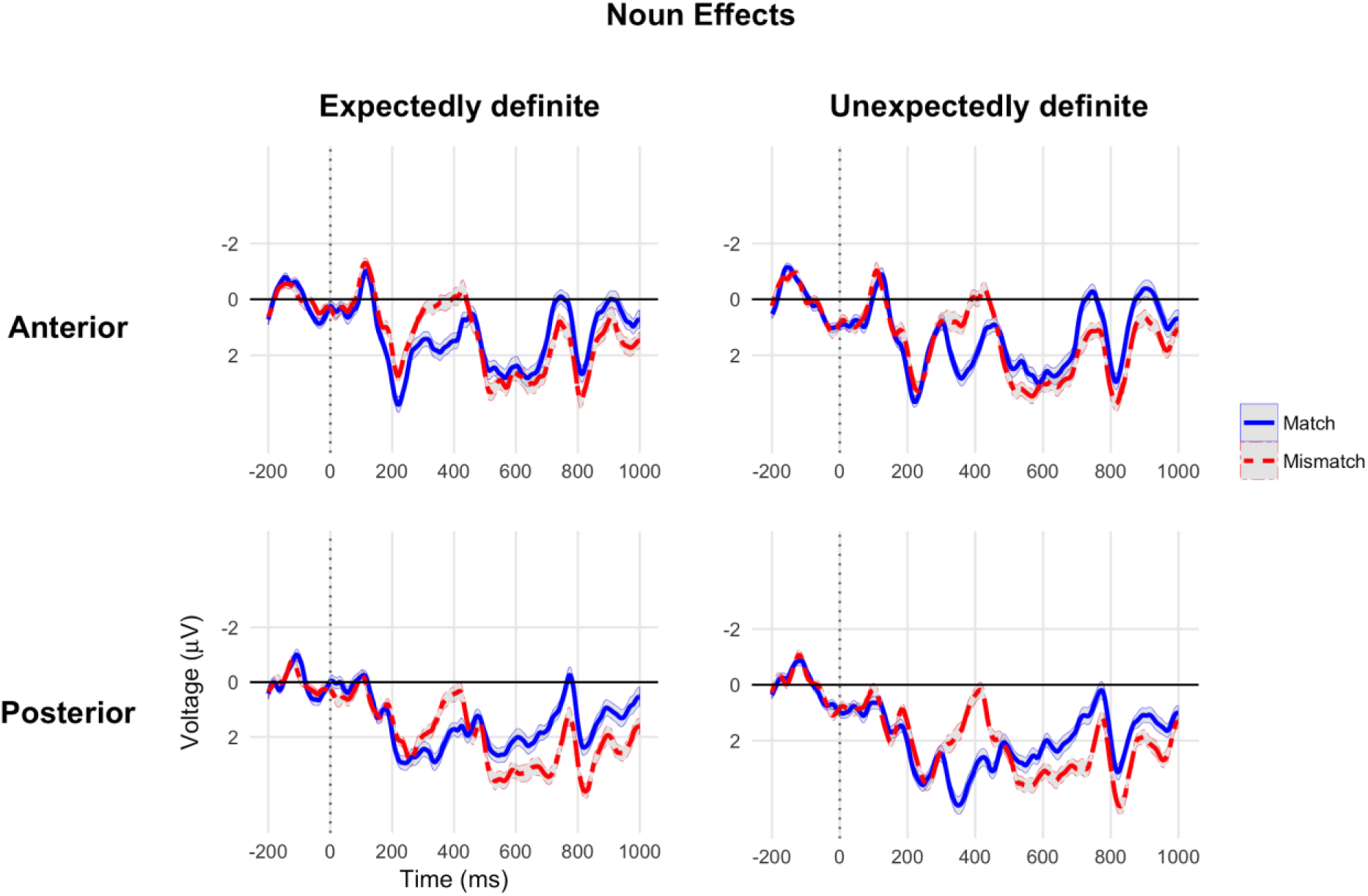
Noun effects. The graphs show the grand-average ERPs elicited by prediction-matching nouns (solid blue lines) and prediction-mismatching nouns (dotted red lines) at the anterior and posterior ROIs (top and bottom graphs, respectively), following articles that were expectedly and unexpectedly definite (left and right graphs, respectively).

**Figure 4.**
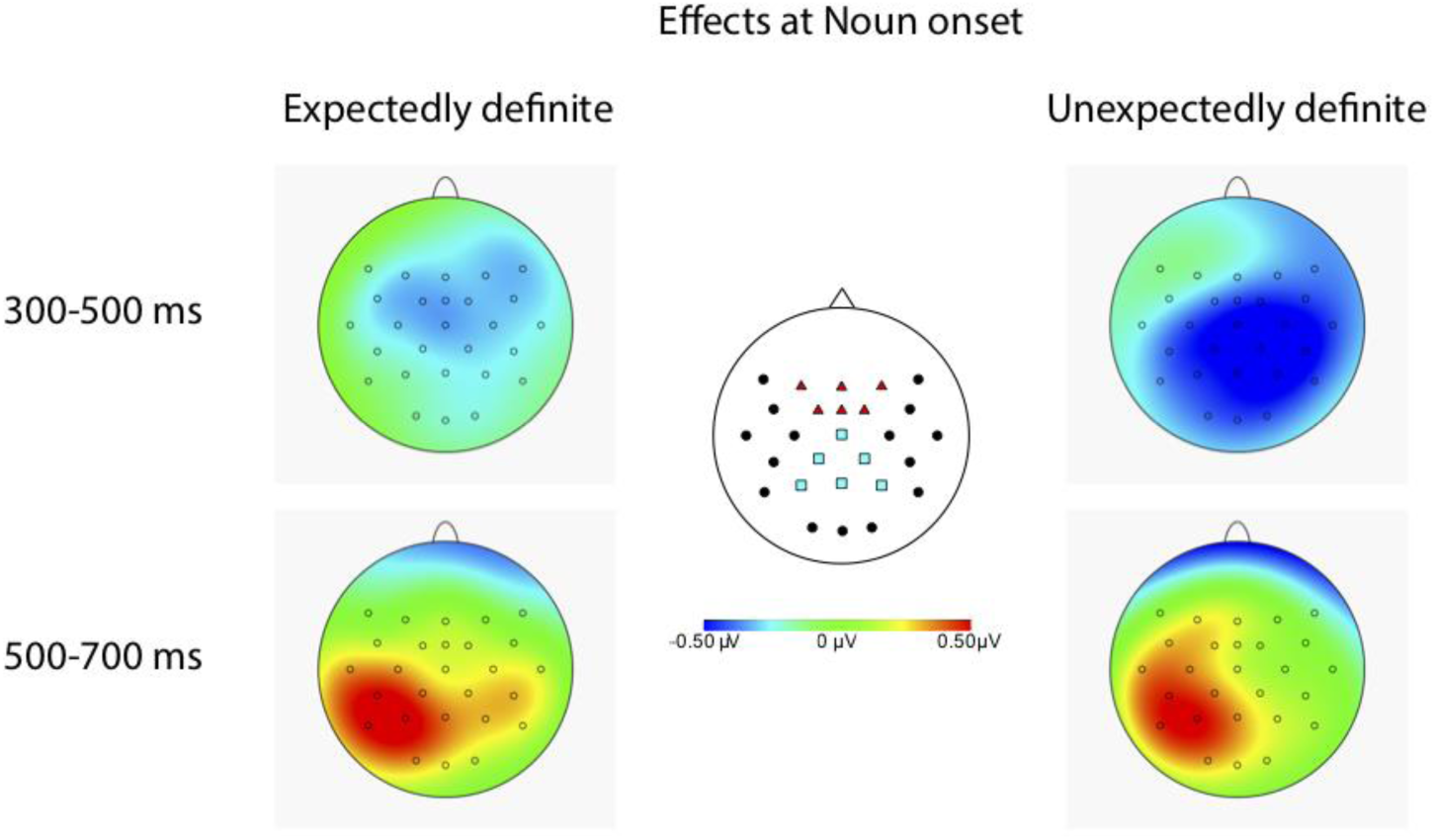
Scalp plots of the noun effects (prediction mismatch minus match) following expectedly and unexpectedly definite articles in both time windows of analysis.

### Exploratory article-analyses

Unlike previous research (Kochari & Flecken, 2018; Otten & Van Berkum, 2009), our pre-registered analyses explicitly controlled for the general effect of article form (‘de/het’), but like previous research our analyses did not take into account potential interactions between article form and the effects of interest. Therefore, we repeated the article analysis with a three-way interaction term between ‘article-form’ (de/het), ‘gender’ (expected/unexpected) and ‘definiteness’ (expected/unexpected); the results of these analyses are listed in the appendix, we only briefly summarize them here. Cloze values for the two article types per condition are listed in Table 4.

**Table 4.** Cloze values (mean *M* and standard deviation *sd*) for ‘de’ and ‘het’

| Table 4. Cloze values (mean $M$ and standard deviation $sd$) for ‘de’ and ‘het’ | | | | |
| --- | --- | --- | --- | --- |
| Definiteness | Gender | Article | $M$ (%) | $sd$ |
| Expected | Match | de | 94.1 | 6.7 |
|  |  | het | 91.4 | 6.9 |
|  | Mismatch | de | 1.2 | 3.2 |
|  |  | het | 1.4 | 3.5 |
| Unexpected | Match | de | 1.1 | 2.8 |
|  |  | het | 0.5 | 1.8 |
|  | Mismatch | de | 0.9 | 2.1 |
|  |  | het | 1.4 | 3.0 |

The results are similar to those obtained in the pre-registered analysis: both time windows showed more negative ERPs (enhanced N400s) for gender-mismatch compared to gender-match, with the strongest difference at posterior electrodes, and likewise for definiteness mismatch. As before, we did not obtain evidence that allowed us to reject the null-hypothesis that the effect of gender-match was the same for expectedly and unexpectedly definite articles. As for the three-way interaction, while our results did not allow us to reject the null hypothesis that ‘de’ and ‘het’ elicited the same effects, the parameter estimates for the interaction is close to or greater than 1 for each of the 4 spatiotemporal ROIs. The results of exploratory follow-up analyses involving only ‘de’ or ‘het’ articles are also in the appendix (for these analyses, we removed the by-item random slope for ‘gender’). We refrain from theoretical conclusions based on these results, but simply note the gender mismatch effect was numerically greater for ‘het’ than for ‘de’, and that the gender by definiteness interaction term for ‘de’ and ‘het’ went in opposite directions.

### Exploratory Bayesian mixed-effects model analyses

Our pre-registered and exploratory article-analyses yielded significant effects of gender-match and definiteness, but non-significant p-values for the interaction between these factors, indicating a failure to reject the null-hypothesis that gender-mismatching articles elicit the same effect when they are expectedly or unexpectedly definite. To more directly examine the support for/against the null-hypothesis, we performed an exploratory Bayes mixed-effects model analysis using the brms package for R (Bürkner, 2016), which fits Bayesian multilevel models using the Stan programming language (Stan Development Team, 2016). All analysis files are available on our OSF page. We constructed models for the posterior ROI in the 300-500 and 500-700 ms time window after article onset, where the gender-mismatch was strongest. These models included as fixed effects the factors definiteness, gender, article type, and all their interactions, and constituted the most maximal model possible with by-subject random slopes for all fixed effects and their interaction, and by-item random slopes for definiteness and gender and their interaction. We included normally distributed, weakly informative priors for the intercept (mean = 0, SD = 2) and for the effects of gender, definiteness, article type and all interactions (mean = 0, SD = 1), and a prior for the correlations of group-level (‘random’) effects using the LKJ(2) prior (Bürkner, 2017). We ran each model with 10.000 iterations (2000 warm-up) in 4 chains (e.g., Vasishth, Beckman, Nicenboim, Li & Kong, 2018; https://rdrr.io/cran/brms/man/bayes_factor.brmsfit.html; see also https://mvuorre.github.io/post/2017/bayes-factors-with-brms/). From these models, we calculated Bayes Factors using the Savage-Dickey method (e.g., Wagenmakers, Lodewyckx, Kuriyal & Grasman, 2010). This method calculates the Bayes Factor (BF^7^) as the ratio between the posterior and prior distribution at an effect size of 0μV, and quantifies the obtained evidence for the alternative hypothesis (H_1_) that there is a non-zero effect, or for the null hypothesis (H_0_) that there is no such effect.

In the two time windows (Figure 5), we obtained anecdotal evidence against an interaction between definiteness and gender (300-500: BF_01_ = 2.01; b = 0.26, 95% Credible Interval (CrI) [−0.56 1.06]; 500-700: BF_01_ = 2.47; b = -.06, CrI [−0.86 0.73]), moderate evidence for an effect of gender (300-500: BF_10_ = 3.6; parameter estimate b = -.50, CrI [−.91 −.09]; 500-700: BF_10_ = 5.7; b = −0.56, CrI [−0.99 −0.13]), extreme and strong evidence for an effect of definiteness (300-500: BF_10_ > 100; b = −1.18, CrI [−1.61 −0.75]; 500-700: BF_10_ = 23.8; b = − 0.75, CrI [−1.22 −0.27]), and essentially no evidence for the three-way interaction between definiteness, gender and article type (300-500: BF_10_ = 1.03; b = 0.56, CrI [−0.73 1.84]; 500-700: BF_10_ = 1.15; b = 0.50, CrI [−.83 1.82]). Estimated means per condition and for the gender-mismatch effect across conditions are shown in Figure 6, which shows that the effect of gender-mismatch goes in the same directions for all comparisons, and is somewhat larger for ‘het’ than for ‘de’.

**Figure 5.**
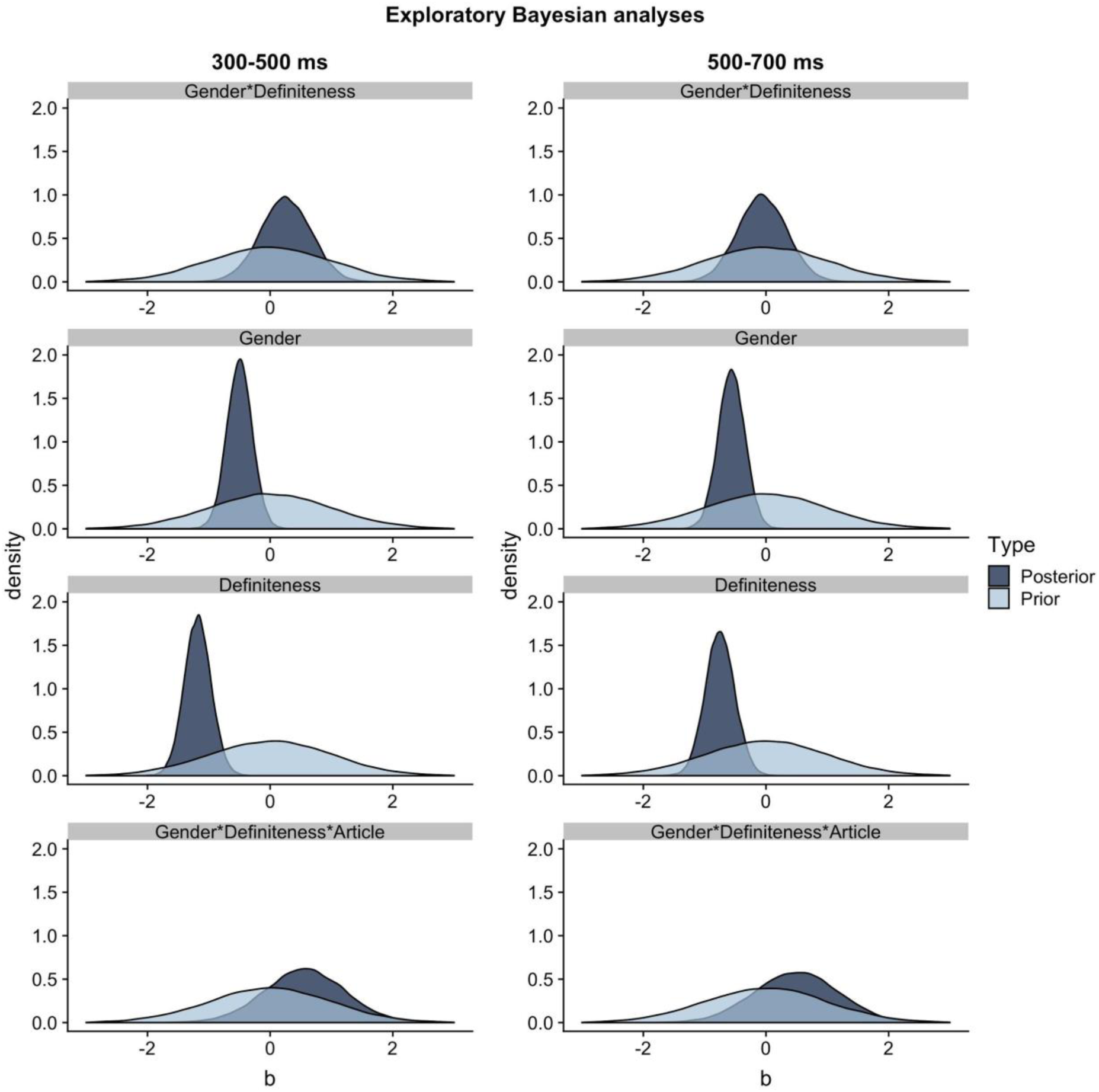
Results of the Bayesian exploratory analyses on data from the posterior ROI in each time window: posterior density for the main effects of and interaction between definiteness and gender (b), all depicted priors have a mean of zero and standard deviation of 1.

**Figure 6.**
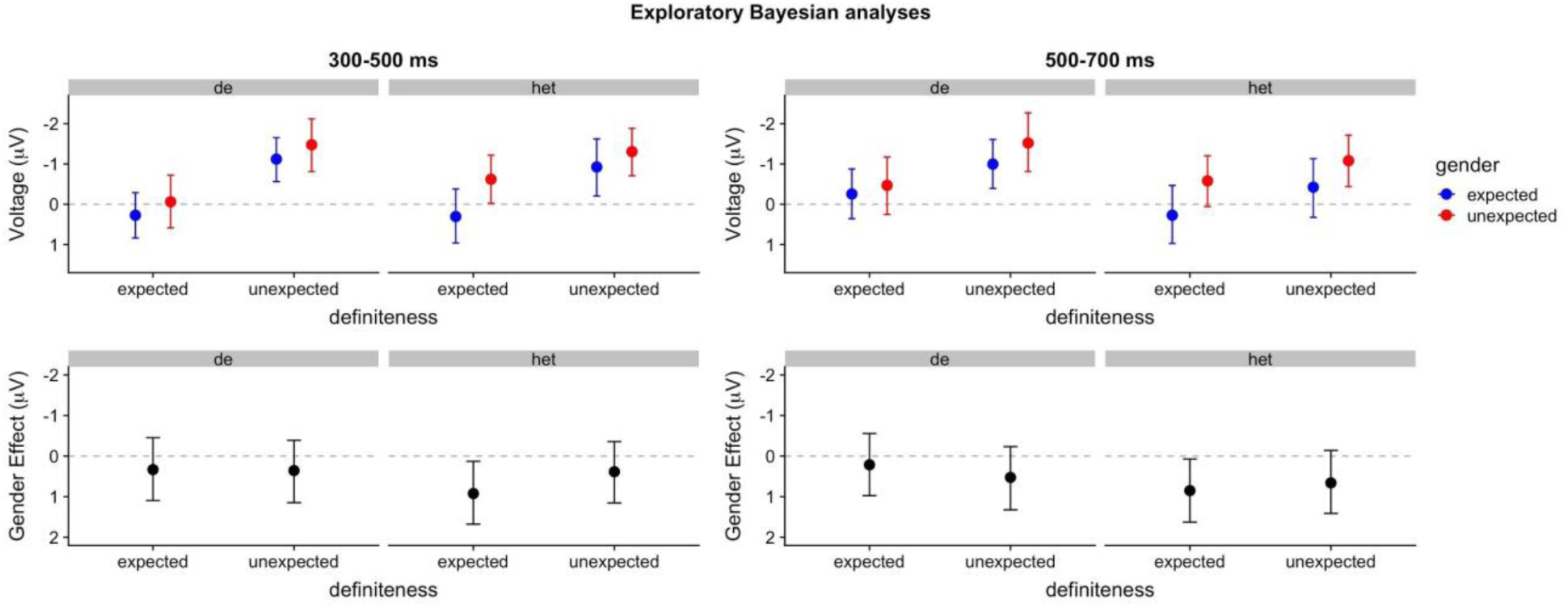
Results of the Bayesian exploratory analyses on data from the posterior ROI in each time window. Top graphs: estimated means and 95% uncertainty interval for each of the conditions. Bottom graphs: estimated means and 95% uncertainty intervals for the effect of gender (mismatch minus match). As in the ERP plots, negative voltage is plotted upwards.

In sum, we conclude from these exploratory analyses^8^ that there is convincing evidence for an effect of gender and definiteness, and that, while there is more evidence against than for their interaction, this evidence is essentially inconclusive.

## DISCUSSION

In this ERP study on Dutch mini-story comprehension, we contrasted two hypotheses about the functional significance of what are known as ‘pre-nominal prediction effects’, the differential neural activity when pre-nominal articles mismatch the gender of a highly predictable noun (Kutas et al., 2011; Van Berkum, 2009), compared to gender-matching articles. One hypothesis is that people predict the article and the noun together (DeLong et al., 2005; Kutas et al., 2011; Wicha et al 2003, 2004; see also Dell & Chang, 2014): people may predict a specific noun including its gender, and then also predict the specific form of the article (which depends on gender). The other hypothesis is that people do not predict the article, but only the noun (with or without its gender). Once they encounter the article, they use its gender-marking to update their prediction of the noun. In both hypotheses, people access the gender of the yet-unseen noun and should thus be sensitive to gender-mismatches, but only in the first hypothesis do people predict the lexical form of the article itself. We contrasted these hypotheses by investigating whether gender effects only occur or are strongest when gender marking is expected (i.e. the article is expectedly definite) versus when gender marking is not expected (i.e. the article is unexpectedly definite). The latter outcome would suggest that prediction of a specific article form is not required to elicit a gender effect.

We created story contexts that strongly suggested either a definite or indefinite noun phrase (e.g., “het/een boek”) as its most predictable continuation, as determined in a cloze probability pre-test. In the ERP experiment, these story contexts were always followed by a definite noun phrase that either matched or mismatched the predictable noun and its gender (‘het boek’ or ‘de roman’, respectively). This allowed us to examine whether a gender effect is observed only when comparing articles that match the predictable gender *and* form to articles that match neither, or whether the gender effect is also observed when both articles have an unexpected form.

Our pre-registered analyses revealed enhanced negativity in the 300-500 ms time window (i.e., increased N400 amplitude) for gender-mismatching articles compared to matching articles, consistent with several previous demonstrations of prediction of specific upcoming words (e.g., Foucart et al., 2014; Martin et al., 2018; Molinaro et al., 2018; Wicha et al., 2003, 2004). As in these previous studies, this effect extended into the subsequent 500-700 ms time window.

Our pre-registered analyses also revealed enhanced N400s for unexpectedly definite articles compared to expectedly definite articles, consistent with a previous report by Schlueter et al. (2018; see also Kirsten et al. 2014). This effect also extended into the subsequent 500-700 ms time window. Unexpected definiteness thus seems to have repercussions for semantic processing. It may cause enhanced semantic processing because it requires a change to the event-based representation of the discourse context (e.g., Clifton, 2013; Frazier, 2006; Zwaan & Radvansky, 1998).

Crucially, although the gender-mismatch effect was numerically larger for expectedly definite articles than for unexpectedly definite articles, we did not obtain convincing evidence for an interaction effect. That is, our results did not allow us to reject the null-hypothesis that the effect of gender-match was similar for expectedly and unexpectedly definite articles. In the 300-500 ms time window, the estimate for the interaction term was about .5 µV but this effect did not reach traditional levels of statistical significance. In the 500-700 ms time window, the estimate for the interaction effect between gender-mismatch and expected definiteness was close to zero. These patterns of results did not change considerably in exploratory analyses wherein effects of gender and/or definiteness could vary for the two article types (‘de/het’). These analyses yielded even weaker interaction between gender and definiteness, but also suggested that ‘de’ and ‘het’ might be associated with different patterns of effects.

Exploratory Bayesian mixed-effect model analyses yielded moderate evidence for an effect of gender, very strong evidence for an effect of definiteness, and some evidence for the lack of an interaction between gender and definiteness. However, the obtained evidence regarding this interaction was merely anecdotal/suggestive and can be considered inconclusive.

### Implications for pre-nominal prediction-effects

Our results do not support previous claims that pre-nominal effects result from article form prediction (DeLong et al., 2005; Kutas et al., 2011; Wicha et al 2003, 2004; see also Dell & Chang, 2014), and could be said to favour an account wherein people only predict the noun (and possibly its gender) and evaluate or update this prediction upon receiving relevant gender information (e.g., Van Berkum et al., 2005). Article form prediction does not seem to be required for the elicitation of pre-nominal prediction effects.

However, our results do not constitute clear evidence *against* article prediction either. We did not obtain convincing evidence in support of the null hypothesis, and the pairwise differences between conditions in the 300-500 ms time window were compatible with an account in terms of article prediction (but as Figure 6 shows, these patterns may be different for the two article types). Maybe our participants did predict article-noun combinations but our study simply failed to find supportive evidence. While that is a possibility, the results in the 500-700 ms time window offered even less evidence that the gender-mismatch effect depended on expected definiteness^9^.

Several further aspects of our results are noteworthy, one being that definiteness elicited a much stronger effect than gender (even if one only considers the gender effect for expectedly definite articles). N400 activity is thus not driven by word predictability (in terms of cloze probability) alone, another indication that the observed effects do not merely reflect detection of prediction mismatch (i.e., mismatch between the predicted and the encountered article), confirming decades of N400 research (Kutas & Federmeier, 2000, 2011). One possible explanation is that our participants pre-activated noun gender *and* definiteness, but pre-activated gender more weakly than definiteness, perhaps because lexical-grammatical features are more specific and ‘harder’ to predict than semantic features (e.g., Pickering & Gambi, 2018). Our results are compatible with such an explanation but do not directly support it. Another explanation, which we tend to prefer, is that, compared to unexpected gender, unexpected definiteness has more meaning and leads to intensified semantic retrieval (e.g., Kutas & Federmeier, 2000, 2011; Van Berkum, 2009) or incurs a greater change to the semantic representation of sentence meaning (Bornkessel-Schlesewsky & Schlesewsky, 2019; Rabovsky, Hansen & McClelland, 2018; Nieuwland et al., 2018). In our experiment, unexpected gender may have signalled a (possibly very small) change in upcoming meaning, for example, instead of ‘church’ participants could update their prediction to a less specific conceptual representation (e.g., some type of building for religious congregation) or some plausible, lexically specific alternative. However, unexpected definiteness has stronger repercussions for the information structure of the discourse, because it is typically reserved for uniquely identifiable referents (ones that are already given, readily accessible or anticipated; Abbott, 2004, 2006; Almor & Nair, 2007; Ariel, 1988; Arnold et al., 2013; Frazier, 2006; Sanford & Garrod, 1988; Schumacher, 2009; Roberts, 2003). Unexpected definiteness therefore violates the presupposition of a uniquely identifiable referent (Karttunen, 1974; Krahmer, 1998; Levinson, 1983; Stalnaker, 1977; Von Fintel, 2004). This could act as a ‘relevance signal’ that triggers more detailed semantic processing or lead to a change in how the meaning of the sentence is represented (by accommodation of a unique referent into the discourse representation, e.g., one specific church; see Beaver, 1999; Von Fintel, 2008). Such changes in semantic processing, and the potential meanings they afforded, may be reflected in N400 activity (Bornkessel-Schlesewsky & Schlesewsky, 2019; Kutas & Federmeier, 2011; Rabovsky et al., 2018; Van Berkum, 2009).

This could be related to a surprising result from our study, namely that predictable nouns elicited smaller N400 amplitude when they followed unexpectedly definite articles compared to expectedly definite articles. The semantic processing changes incurred by unexpectedly definite articles (e.g., intensified semantic retrieval or updating of sentence meaning) could have boosted the semantic pre-activation of the predictable noun. While this could be an interesting novel conclusion, we refrain from such a theoretical claim, because the ERPs elicited by the nouns may be confounded by activity elicited by the articles. For example, the baseline period for the noun analysis happens to be the time period where the N400 effect of unexpected definiteness was most pronounced (400-600 ms after article-onset), and it is unclear whether effects of the articles extended beyond this period and in what manner. Such issues have been known to plague interpretation in other ERP paradigms (for a review, see Steinhauer & Drury, 2012; Nieuwland, 2019), and highlight the need for caution to interpret ERP effects associated with subsequent words.

A final observation worth discussing is that the two Dutch articles (‘de/het’) appeared to elicit different effects (see also Brouwer, Sprenger & Unsworth, 2017; Loerts, Wieling & Schmid, 2013). We acknowledge that our results do not convincingly demonstrate this difference, and more reliable estimates of the interaction patterns are needed. Nevertheless, it is worthwhile to consider why this difference may occur in the first place. Previous studies (Kochari & Flecken, 2018; Otten & Van Berkum, 2009) counterbalanced the expected form of the article (‘de/het’) across items, but did not report effects of the articles themselves, so previous literature did not allow for a strong a priori hypothesis. One relevant factor could be the high frequency of ‘de’ (common-gender) compared to ‘het’ (neuter-gender), and the fact that people are faster at article assignment and verification for ‘de’ words than for ‘het’ words (Deutsch & Wijnen, 1985). If pre-nominal ERP effects reflect the detection of a mismatch between predicted and expected gender, and gender-mismatch is detected more easily or faster for ‘de’ than ‘het’, then ‘de’ should have elicited a larger effect than ‘het’. However, this is not what we found, and we do not believe that N400 amplitude reflects a prediction mismatch detection process (see also Kutas & Federmeier, 2011; Rabovsky et al. 2018; Szewczyk & Wodniecka, 2018; Van Berkum, 2009). Alternatively, unexpected ‘het’ may elicit a larger gender effect because ‘het’ is a lower-frequent word taking the place of the more frequent article ‘de’. Unexpected ‘het’ therefore possibly allows for fewer alternative plausible noun continuations than unexpected ‘de’ (e.g., Brouwer et al., 2017). Yet another potentially relevant factor is that ‘de’ and ‘het’ are not perfectly diagnostic of noun gender: ‘de’ is also the plural definite article irrespective of noun gender, and ‘het’ is always used for diminutive nouns irrespective of gender. It is unknown, however, whether such differences matter in an experiment where participants do not encounter plural or diminutive continuations.

Importantly, the potential processing differences associated with ‘de’ and ‘het’ highlight the relevance of cross-linguistic differences in studies on linguistic prediction (see also Bornkessel-Schlesewsky & Schlesewsky, 2009; Kamide, Scheepers, & Altmann, 2003; Van Bergen & Flecken, 2017). For example, Dutch does not mark gender on indefinite articles and its article form for definites is not a perfectly reliable cue to noun gender, but this is different in Spanish, which uses a different and unique article for each possible combination of gender, definiteness and number. Different patterns of results may therefore be expected for Spanish: for example, stronger evidence for article form prediction, and weaker evidence for processing differences associated with different genders. Yet different patterns of results may occur in languages with rich case and gender systems such as German or Polish.

### Conclusions

Our results add to the growing body of evidence for linguistic prediction using a pre-nominal gender manipulation in highly constrained sentences. We did not obtain clear evidence that the effect of gender-mismatch depended on whether the presence of gender marking was expected given the context. Although these results do not constitute clear evidence *against* prediction of a specific article form, they suggest that article form prediction is not *required* to elicit a pre-nominal effect.

## Appendix

| Exploratory analysis for article-elicited ERPs, including the 3-way interaction. |  |  |  |  |  |  |  |
| --- | --- | --- | --- | --- | --- | --- | --- |
|  |  | Time window |  |  |  |  |  |
| Factor | ROI | 300-500 ms<br>(N400) |  |  | 500-700 ms<br>(post-N400) |  |  |
| | | $\beta$, $CI$ | $t$ | $p$ | $\beta$, $CI$ | $t$ | $p$ |
| Gender | Anterior | 0.11,<br>[-0.49, 0.27] | -0.55 | 0.58 | -0.32,<br>[-0.73, 0.09] | -1.55 | 0.12 |
|  | Posterior | -0.52,<br>[-0.89, -0.15] | -2.78 | 0.00 | -0.59,<br>[-0.99, -0.20] | -2.93 | 0.00 |
| Definiteness | Anterior | -1.15,<br>[-1.53, -0.76] | -5.87 | <0.001 | -0.56,<br>[-0.96, -0.15] | -2.66 | 0.001 |
|  | Posterior | -1.25,<br>[-1.62, -0.88] | -6.64 | <0.001 | -0.83,<br>[-1.23, -0.43] | -4.10 | <0.001 |
| Gender:Defini<br>teness | Anterior | -0.08,<br>[-0.85, 0.69] | -0.20 | 0.84 | -0.57,<br>[-0.88, 0.76] | -0.14 | 0.89 |
|  | Posterior | 0.28,<br>[-0.46, 1.02] | 0.74 | 0.46 | -0.09,<br>[-0.89, 0.71] | -0.22 | 0.83 |
| Gender:Defini<br>teness:Article | Anterior | 1.51,<br>[-0.02, 3.04] | 1.94 | 0.05 | 1.01,<br>[-0.63, 2.64] | 1.20 | 0.23 |
|  | Posterior | 1.01<br>[-0.47, 2.48] | 1.34 | 0.18 | 1.00,<br>[-0.59, 2.59] | 1.23 | 0.22 |

| Follow-up analysis for articles in the 300-500 ms time window (N400) |  |  |  |  |  |  |  |
| --- | --- | --- | --- | --- | --- | --- | --- |
|  |  | Article |  |  |  |  |  |
| Factor | ROI | de |  |  | het |  |  |
| | | $\beta$, $CI$ | $t$ | $p$ | $\beta$, $CI$ | $t$ | $p$ |
| Gender | Anterior | 0.05,<br>[-0.57, 0.67] | 0.16 | 0.87 | -0.26,<br>[-0.90, 0.38] | -0.80 | 0.42 |
|  | Posterior | -0.33,<br>[-0.89, 0.23] | -1.16 | 0.25 | -0.71,<br>[-1.30, -0.11] | -2.34 | 0.02 |
| Definiteness | Anterior | -1.21,<br>[-1.75, -0.67] | -4.41 | <0.001 | -1.04,<br>[-1.59, -0.48] | -3.68 | <0.001 |
|  | Posterior | -1.54,<br>[-2.06, -1.02] | -5.80 | <0.01 | -1.01,<br>[-1.54, -0.48] | -3.73 | <0.001 |
| Gender:Defini<br>teness | Anterior | -0.81,<br>[-1.88, 0.26] | -1.48 | 0.14 | 0.69,<br>[-0.40, 1.78] | 1.24 | 0.21 |
|  | Posterior | -0.22,<br>[-1.26, 0.81] | -0.43 | 0.67 | 0.81,<br>[-0.24, 1.86] | 1.51 | 0.13 |

| Follow-up analysis for articles in the 500-700 ms time window ( post-N400) |  |  |  |  |  |  |  |
| --- | --- | --- | --- | --- | --- | --- | --- |
|  |  | Article |  |  |  |  |  |
| Factor | ROI | de |  |  | het |  |  |
| | | $\beta$, $CI$ | $t$ | $p$ | $\beta$, $CI$ | $t$ | $p$ |
| Gender | Anterior | -0.22,<br>[-0.85, 0.42] | -0.67 | 0.50 | -0.43,<br>[-1.06, 0.19] | -1.36 | 0.18 |
|  | Posterior | 0.37,<br>[-0.98, 0.25] | -1.17 | 0.24 | -0.82,<br>[-1.40, -0.24] | -2.78 | 0.01 |
| Definiteness | Anterior | -0.52,<br>[-1.09, 0.06] | -1.76 | 0.08 | -0.53,<br>[-1.12, 0.06] | -1.75 | 0.08 |
|  | Posterior | -1.00,<br>[-1.56, -0.44] | -3.52 | <0.001 | -0.63,<br>[-1.21, -0.06] | -2.16 | 0.03 |
| Gender:Defini<br>teness | Anterior | -0.54,<br>[-1.68, 0.60] | -0.92 | 0.36 | 0.45,<br>[-0.72, 1.62] | 0.75 | 0.45 |
|  | Posterior | -0.57,<br>[-1.68, 0.53] | -1.01 | 0.31 | 0.41,<br>[-0.72, 1.55] | 0.71 | 0.48 |

## Footnotes

1 The reported effects do seem to differ, at least visually, from typical N400 effects elicited with predictable vs unpredictable *nouns* with respect to latency and scalp distribution.

2 A parallel can be drawn to the literature on activation of gender information during word production. Some models of production argue that gender information is activated when people access a specific word meaning (lexical access), but other models argue that gender information is only activated when people access a phonological form (since that form may depend on gender; for discussion, see Caramazza, 1997; Roelofs, Meyer & Levelt, 1998; Schiller & Caramazza, 2006; Schriefers & Jescheniak, 1999).

3 We were unable to obtain the raw cloze responses from Otten & Van Berkum (2009). In Kochari & Flecken (2018), which used materials based on the Otten and Van Berkum study, 45% of all cloze responses contained an indefinite article (97 out of the total number of 112 items contained at least one response with an indefinite article, and in 62 out of all items, more than half the responses contained the indefinite ‘een’).

4 In both studies, the gender-match and -mismatch conditions start to diverge as early as 0 ms after article onset, and continue to diverge into the later time windows. Given that an effect as early as that is psychologically implausible, an alternative explanation is that these effects reflect a slow signal drift associated with voltage differences in the baseline period. In other words, it is not clear to what extent the obtained effects are truly generated by the article (see also Ito et al., 2017a,c; Nieuwland et al., 2018), and whether the obtained effects would hold when a countermeasure is performed to deal with the potential baseline problem (e.g., applying a 0.1 Hz filter instead of the 0.03 Hz filter, or applying a post-onset baseline correction; see also Ito et al., 2017b).

5 Our pre-registration was submitted after having tested five participants, and included our target sample size (N=48), experimental design and stimulus list assignment, data pre-processing, data exclusion and statistical analysis on AsPredicted.org via OSF (https://osf.io/mv4hw/). Analyses that were not pre-registered are labelled as exploratory.

6 Our pre-registered analyses in the 500-700 ms time windows converged but revealed random effect correlations of ±1, indicating overfitting. Re-running these analyses after removing the relevant random slope did not meaningfully change the observed pattern of effects.

7 We follow Wetzels and Wagenmakers (2012) in evidence-categorization of the obtained Bayes Factors (see also Jeffreys, 1961), but note that the obtained values themselves are meaningful: BF_10_ = 1, in which the subscript refers to the evidence favouring the alternative hypothesis over the null-hypothesis (=1/BF_01_), means that the data are precisely as likely to occur under the alternative hypothesis as the null hypothesis, BF_10_ = 3 (or BF_01_ = ⅓) means that the data 3 times more likely to occur under the alternative hypothesis than the null hypothesis, whereas BF_01_ = 3 (or BF_10_ = ⅓) means the data are 3 times more likely under the null hypothesis than the alternative hypothesis.

8 We emphasize that the Bayes Factors (much more so than the parameters estimates) are influenced by the choice of priors. A zero-mean prior with SD = 1 means that 95% of the plausible values lie within −2 to 2 µV. Using a prior with a smaller standard deviation around zero (e.g., SD = .5) increases the prior density at an effect size of zero (i.e., increasing confidence in a null-effect) and makes it harder to obtain evidence for the null hypothesis. Consequently, such a prior weakens the anecdotal evidence against an interaction between definiteness and gender (300-500: BF_01_ = 1.29; 500-700: BF_01_ = 1.51), while somewhat strengthening or having little impact on the evidence for an effect of gender, definiteness, or the 3-way interaction. Alternatively, one could change the mean prior effect size of the effects of gender, definiteness and the interaction to .5 µV, compatible with the effect of gender being driven entirely by the expectedly definite articles. This prior hardly changes the evidence for the null hypothesis in the 300-500 ms time window (BF_01_ = 1.98) but somewhat strengthens this evidence in the later window (BF_01_ = 2.74).

9 It is possible that these two time windows are associated with different processes. The earlier time window might be dominated by semantic activation processes whereas the later one might be dominated by integrative processes (see Nieuwland et al., 2019, for results that are compatible with such a temporal distinction). The two hypotheses contrasted in this study may therefore both be correct.

